# Interplay between consensus and divergent RNA polymerase II C-terminal domain repeats in viability and targeting

**DOI:** 10.1101/448506

**Authors:** Feiyue Lu, Bede Portz, David S. Gilmour

## Abstract

The carboxy-terminal domain (CTD) of RNA polymerase II (Pol II) is composed of repeats of the consensus YSPTSPS, and is an essential binding scaffold for transcription-associated factors. Metazoan CTDs have well-conserved lengths and sequence compositions arising from the evolution of divergent motifs, features thought to be essential for development. To the contrary, we show that a truncated CTD composed solely of YSPTSPS repeats fully supports *Drosophila* viability, but a CTD with enough YSPTSPS repeats to match the length of the wild-type *Drosophila* CTD is defective. Furthermore, a fluorescently-tagged CTD lacking the rest of Pol II dynamically enters transcription compartments, indicating that the CTD functions as a signal sequence. However, CTDs with too many YSPTSPS repeats are more prone to localize to static nuclear foci independent of the chromosomes. We propose that the sequence complexity of the CTD offsets aberrant behavior caused by excessive repetitive sequences without compromising its targeting function.

## Introduction

Transcription in eukaryotes involves coordination of RNA synthesis, RNA processing and modulation of chromatin structure. The CTD of the largest Pol II subunit, Rpb1, plays a central role by serving as a landing pad for many of the proteins involved in these processes (Corden, 2013; Egloff et al., 2012; Eick and Geyer, 2013; Harlen and Churchman, 2017; Zaborowska et al., 2016). There is considerable variation in the amino acid sequence composition of CTDs across different evolutionary lineages (Yang and Stiller, 2014). For example, the yeast CTD and the proximal half of the mammalian CTD are composed primarily of the repeating consensus heptad, YSPTSPS, whereas the distal half of the mammalian CTD consists primarily of divergent motifs, which differ from the consensus at one or more positions (Figure 1A). In addition, the length of the CTD roughly scales with evolutionary complexity (Yang and Stiller, 2014). The divergent motifs and CTD length are postulated to have evolved to provide additional layers of control for gene expression programs essential for development of multicellular organisms (Chapman et al., 2008; Corden, 2013; Dias et al., 2015; Egloff et al., 2012; Eick and Geyer, 2013; Schröder et al., 2013; Simonti et al., 2015; Sims et al., 2011; Zaborowska et al., 2016; Zhao et al., 2016), although this remains largely untested.

**Figure 1.**
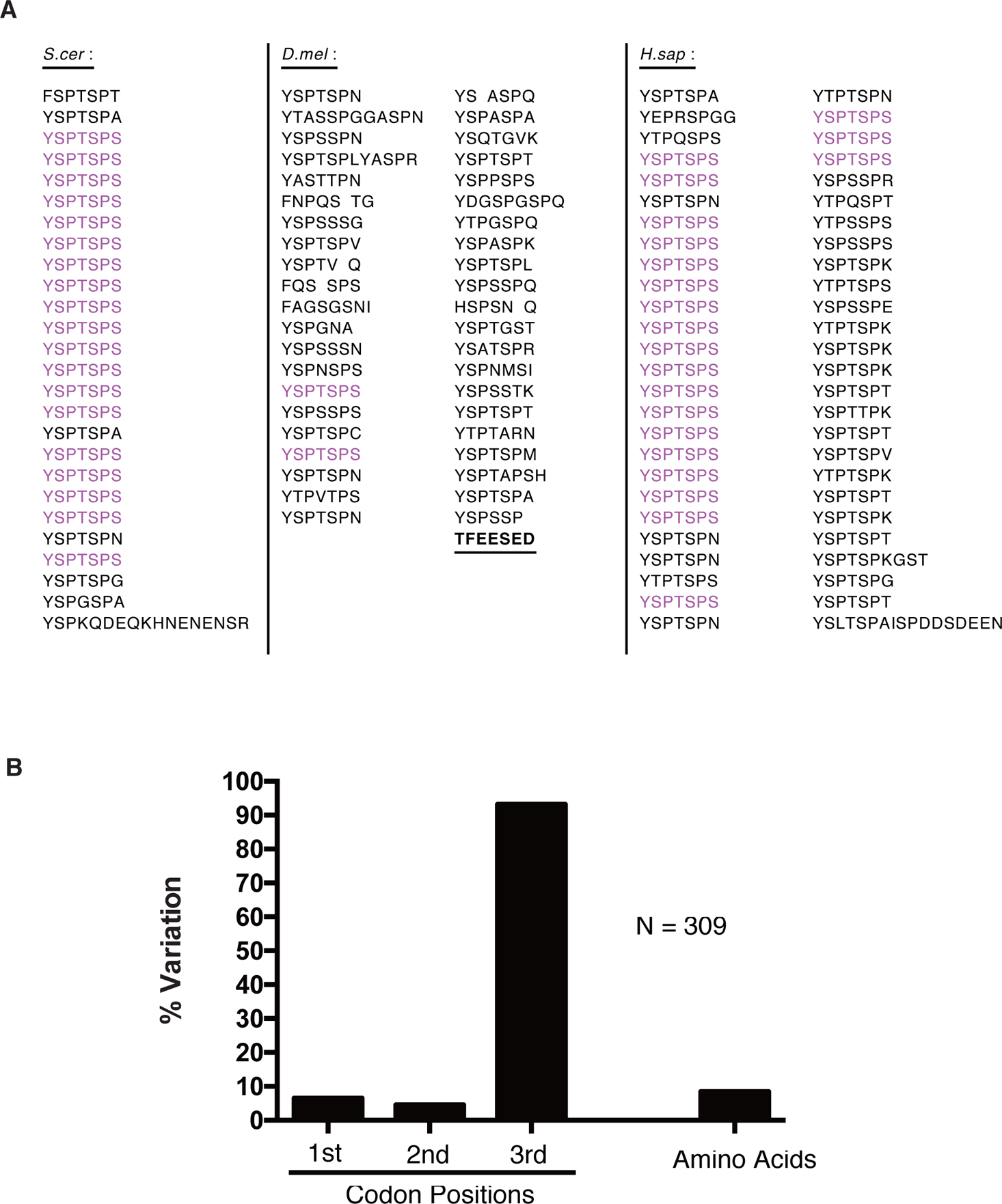
The CTD of *Drosophila* is composed primarily of divergent motifs that are under selective pressure. (A) Display of the CTD of *S. cerevisiae*(26 repeats), *D. melanogaster*(42 repeats) and *H. sapien* (52 repeats) with consensus heptads highlighted in magenta. The acidic tip (underlined) was retained in all consensus CTD mutants described throughout this study. (B) Variation in the codon positions and amino acids of the *Drosophila* CTD. Variation was defined as having at least one sequence variant among 12 species of *Drosophila*.

Here, we assess the significance of sequence complexity and length of the CTD in animal development and viability by using *Drosophila* as a model. The *Drosophila* CTD contains 42 repeats, only 2 of which match the consensus heptad (Figure 1A). Surprisingly, we find that CTDs composed solely of YSPTSPS repeats are sufficient to support fly development. In addition, although consensus CTDs at roughly half wild-type lengths fully support development, consensus CTDs at wild-type lengths do not. This indicates that having too many consensus heptads is deleterious, which can be offset by either shortening the CTD or substituting consensus heptads with divergent motifs. These results argue against the notion that highly-conserved, lineage-specific divergent motifs and the conserved length of the CTD are essential for metazoans. Finally, we show that a GFP-CTD fusion dynamically enters transcription compartments. This defines the CTD as a signal sequence that aids in targeting Pol II to sites of active transcription. Furthermore, the nuclear distribution of the CTD is altered upon substitutions with consensus CTDs in manner that explains the behavior of the Pol II CTD mutants. A GFP-CTD fusion harboring a truncated CTD that fails to support viability in the context of Pol II is depleted from sites of active transcription relative to the wild-type CTD. Likewise, CTDs harboring too many consensus motifs form static nuclear foci when fused to GFP, and fail to support viability in the context of Pol II. Our results suggest the sequence of the CTD has evolved to balance length and composition in order to dynamically target Pol II to transcription compartments without resulting in aggregation.

## Results

### A CTD Solely Composed of 29 Repeats of Consensus Heptads Is Sufficient for Viability of *Drosophila*

Differences between the amino acid sequence conservation of the CTD and underlying nucleic acid sequence for *Drosophila* species indicates that the CTD is subject to significant selective pressure (Figure 1B and Figure S1). Leveraging the sequence complexity of the *Drosophila* CTD, we set out to test the long-standing hypothesis that the divergent motifs enriched among metazoan CTDs and conserved within lineages serve crucial roles in development. Previously, we found that ectopic expression of an RNAi-resistant Rpb1 mutant with as few as 29 *Drosophila* motifs, including the two consensus heptads, rescued lethality caused by depleting endogenous Rpb1 with RNAi (Gibbs et al., 2017). To dissect whether viability was conferred by the remaining divergent motifs, we matched the length of this mutant by replacing the *Drosophila* CTD with a FLAG-tagged, all-consensus CTD (Rpb1^29con^ hereafter) composed of 29 YSPTSPS heptads. Following CRISPR-Cas9 replacement of sequences encoding the wild-type CTD, we obtained homozygous viable flies whose only Rpb1 contained 29 consensus heptads (Figure 2A). For comparison, two fly lines with either an HA tag or a FLAG tag appended to the C-terminus of a wild-type *Drosophila* CTD were generated (Rpb1_HA_ and Rpb1_FLAG_ respectively hereafter). *Rpb1^29con^* flies were indistinguishable from these wild-type counterparts. They showed similar hatch rates and percentages of individuals that completed development when grown at 24^°^C (Figure 2B and 2C). When raised at 30^°^C, these three fly lines were also morphologically indistinguishable, showed similar hatch rates and similar percentages of individuals developing into pupae (Figure 2D and 2E). In addition, *Rpb1^29con^* flies recovered normally after cold shock (Figure 2F). *Rpb1^29con^* flies were mated to *Rpb1_HA_* flies to generate larvae expressing both forms of Pol II. Immunostaining of polytene chromosomes for the two forms of Rpb1 with anti-FLAG and anti-HA antibodies revealed nearly identical staining patterns both before and after heat shock (Figure 2G). In contrast to the hypothesized role of divergent motifs in metazoa, these observations show that Pol II harboring a fully-consensus CTD is sufficient to execute the complex gene regulatory programs necessary for development, reproduction and thermotolerance.

**Figure 2.**
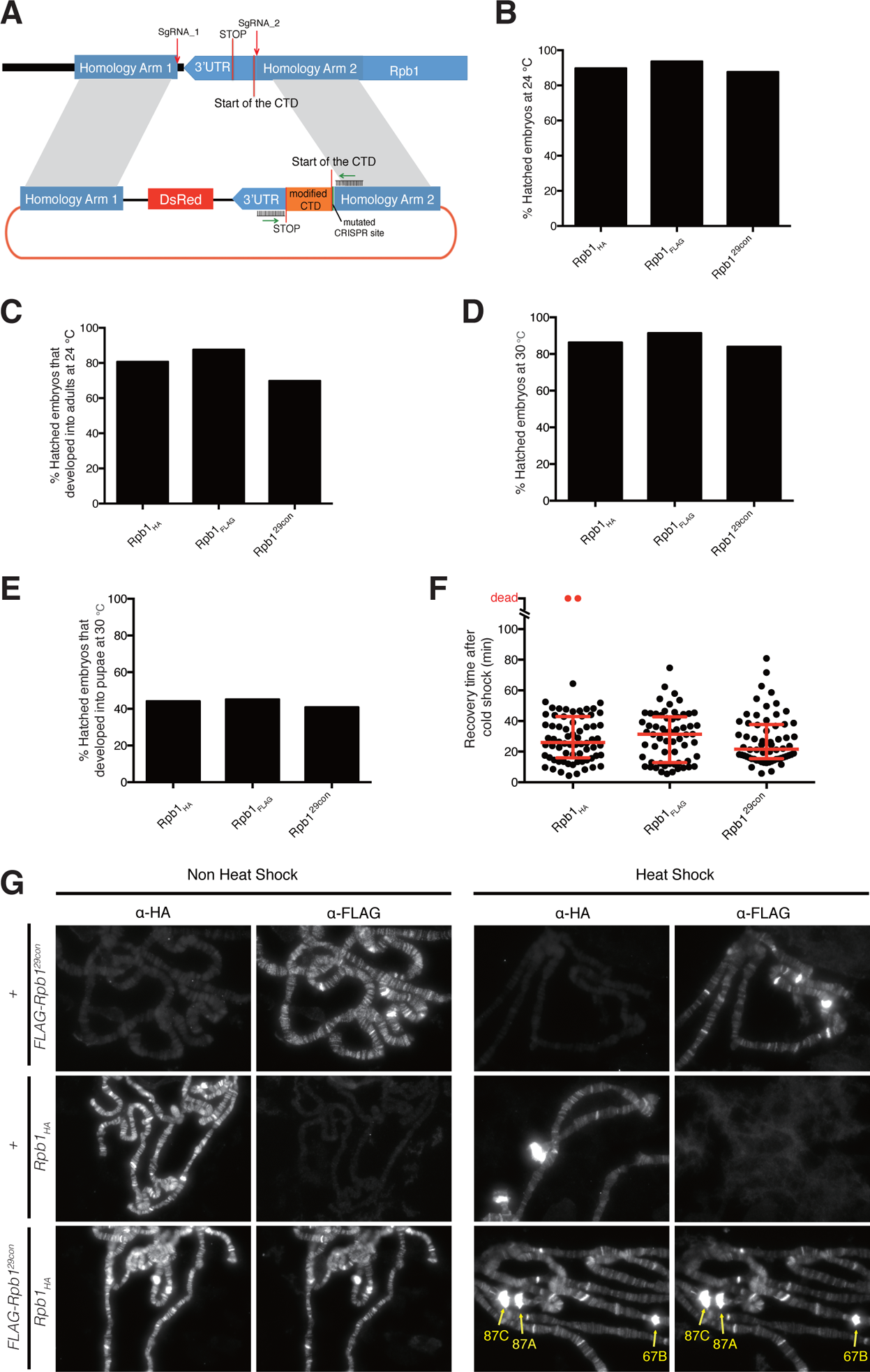
Replacing endogenous Rpb1 with one that has 29 consensus heptads produces healthy homozygous flies. (A) Schematic for CRISPR-mediated replacement of the endogenous *Drosophila* CTD. The region containing each modification (orange) was PCR amplified from genomic DNA using primers flanking the CTD (green arrows) and sequenced. Rpb1_HA_ and Rpb1_FLAG_ have the wild-type *Drosophila* CTD sequence with a HA or FLAG-tag attached to the end of the C-terminus. Rpb1^29con^ has 29 YSPTSPS motifs substituted for the wild-type CTD and a FLAG-tag attached to the end of the C-terminus. (B) Hatch rates of embryos at 24^°^C, measured 36h after egg deposition, n>300 for each genotype. (C) Percentages of hatched embryos that developed into adults when raised at 24^°^C, n>200 for each genotype. (D) Hatch rates of embryos at 30^°^C, measured 36h after egg deposition, n>300 for each genotype. (E) Percentages of hatched embryos that developed into pupae when raised at 30^°^C, n>200 for each genotype. (F) Quantification of recovery time after 14h of cold shock on ice, n=70 for *Rpb1_HA_*; n=64 for *Rpb1_FLAG_*; n=64 for *Rpb1^29con^* (equal numbers of male and female adults were assayed). Dots represent individual adults. Red bars show the medians and interquartile ranges. (G) Immunofluorescence with anti-HA and anti-FLAG antibodies on polytene chromosomes derived from salivary glands of third-instar larvae (FLAG-Rpb1^29con^/Rpb1_HA_) under non heat shock and heat shock conditions. Chromosomes derived from salivary glands lacking either tagged form of Rpb1 (*FLAG-Rpb1^29con^/+* or *Rpb1_HA_/+*) served as negative controls. The heat shock puffs are readily visible as pointed by yellow arrows.

### A Consensus CTD Matching the Wild-type Length of the *Drosophila* CTD Is Defective

Like sequence complexity, the length of the CTD is highly conserved within taxonomic lineages, and is also thought to be essential for viability (Chapman et al., 2008; Corden, 2013; Eick and Geyer, 2013; Liu et al., 2010; Yang and Stiller, 2014; Zaborowska et al., 2016). However, the role of conserved CTD length has been difficult to study in developmentally complex organisms, where changing CTD length also alters motif composition. Our finding that Pol II with a fully-consensus CTD functions in the fly enabled us to directly test the role of altered CTD length on fly viability, divorced from the confounding influence of altered numbers of divergent motifs. To this end, we used our previously described RNAi rescue assay to test if CTDs with 10, 20, 24, 42 or 52 consensus heptads (Rpb1^10con^, Rpb1^20con^, Rpb1^24con^, Rpb1^42con^ and Rpb1^52con^) could restore viability in the presence of lethal levels of Rpb1 knockdown (Gibbs et al., 2017; Portz et al., 2017). Expression of RNAi-resistant forms of Rpb1^20con^ or Rpb1^24con^ restored viability caused by ubiquitous Rpb1 knockdown, whereas expression of Rpb1^10con^ or Rpb1^52con^ did not (Figure 3A and 3B). In contrast to Rpb1^52con^, the human CTD that also has 52 repeats (Rpb1^human^) was able to restore viability (Portz et al., 2017), raising the possibility that the divergent motifs counteract defects caused by having too many consensus heptads. Likewise, the Rpb1^42con^, a mutant closely matching the length of the *Drosophila* CTD, restored viability into adulthood but was markedly less efficient than Rpb1^WT^ (Figure 3B and Figure S2). Importantly, all Rpb1 derivatives were expressed at comparable levels and endogenous Rpb1 was efficiently depleted (Figure 3C). Therefore, the differences in rescue efficiency was not due to variation in the level of expression or variation in the level of Rpb1 knock down. We also performed a wing-specific knockdown of the endogenous Rpb1 and tested the effects of co-expressing various forms of mutant Rpb1. The wing-specific knockdown of Rpb1 gave rise to severely distorted, miniature-sized wings that were rescued by the expression of Rpb1^WT^, Rpb1^20con^, Rpb1^24con^ and Rpb1^29con^. As was observed in the whole-animal experiments, expression of Rpb1^10con^ failed to rescue wing defects. Conversely, expression of either Rpb1^42con^ or Rpb1^52con^ in the wing partially rescued the RNAi phenotype (Figure 3D). This suggests that the basis for the defects caused by reducing the number of consensus heptads may be different from defects caused by having too many consensus heptads.

**Figure 3.**
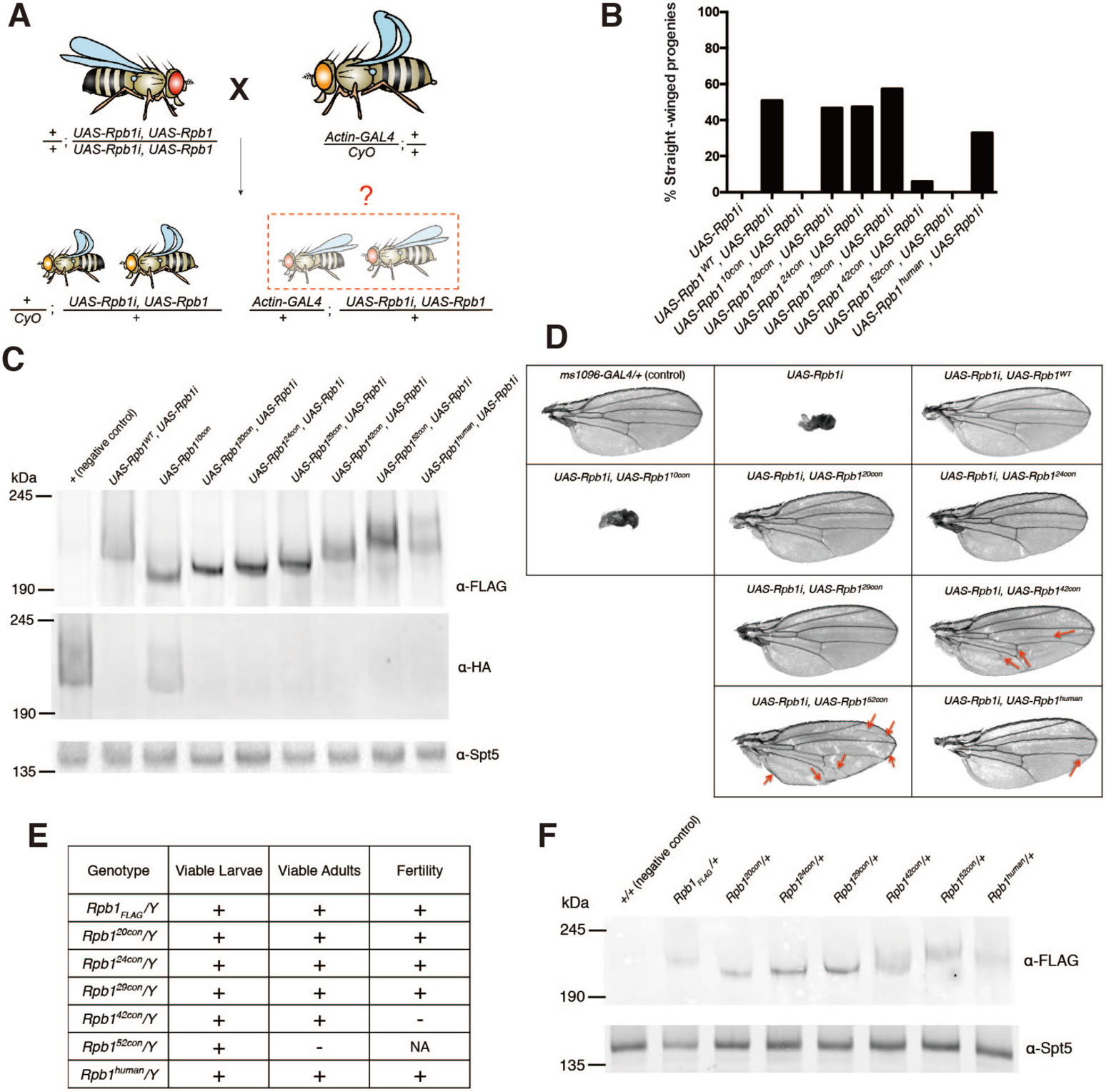
CTDs with too few or too many consensus heptads are dysfunctional. (A) Schematic of the ubiquitous Rpb1i rescue assay. The *Act-GAL4* transgene expresses GAL4 throughout development and drives expression of UAS-associated transgenes. *UAS-Rpb1i* is a GAL4-activated transgene encoding RNAi that knocks down expression of endogenous Rpb1. Rpb1i expression in the absence of ectopically expressed Rpb1 eliminates all straight-winged adult progeny (*Act-GAL4/+; UASRpb1i/+*). Co-expression of a functional derivative of Rpb1 that has been rendered resistant to the RNAi by synonymous mutations rescues the lethality and yields straight-winged progeny. (B) Quantification of the results from the Rpb1i rescue assay, n>150 for each genotype. 10con, 20con, 24con, 29con, 42con, and 52con designate Rpb1 derivatives with 10 to 52 consensus heptads. WT designates the wild-type *Drosophila* CTD and human designates the human CTD. (C) Western blots of late pupae showing the expression of FLAG-tagged, RNAi-resistant forms of Rpb1 derivatives, and the knockdown of the RNAi-sensitive *Rpb1_HA_*. Spt5 serves as a loading control. *UAS-Rpb1* and *Act-GAL4* were each marked with *w+*, thus allowing late pupae with *Act-GAL4* to be distinguished from their *CyO* counterparts by an additional copy of *w+*. (D) Testing the function of various Rpb1 derivatives in *Drosophila* wings. Wing-specific expression was driven by *ms1096-GAL4*. Red arrows indicate creases in the wing or changes in the vein path. (E) Functionality of CRISPR derivatives of Rpb1 in males. The *Rpb1* gene resides on the X chromosome. (F) Western blot showing the expression of FLAG-tagged, CRISPR derivatives of Rpb1 in late pupae. Spt5 serves as a loading control.

To definitively test if the simplified CTDs can replace the functions of the wild-type CTD, we used CRISPR-Cas9 to introduce various mutations into the endogenous Rpb1 locus in flies. *Rpb1^20con^*, *Rpb1^24con^*, *Rpb1^42con^* and *Rpb1^52con^* were introduced whereas two attempts to introduce Rpb1^10con^ were unsuccessful. As was the case for *Rpb1^29con^*, *Rpb1^20con^* and *Rpb1^24con^* supported development and proliferation (Figure 3E, Figure S3A and S3B, Figure S4). They also functioned like *Rpb1_FLAG_* under normal growth conditions and in response to stress (Figure S3C to S3E). Hemizygous males carrying Rpb1^42con^ were sterile whereas Rpb1^52con^ was unable to support development into adulthood. In contrast to Rpb1^52con^, Rpb1^human^ substituted for the endogenous Rpb1 to produce homozygous viable adults that were fertile (Figure 3E, Figure S4). The deficiency of Rpb1^42con^ and Rpb1^52con^ was not simply due to under-expression, as all CRISPR derivatives of Rpb1 were expressed at comparable levels (Figure 3F). Collectively, these results suggest that while a fly can survive without divergent motifs, too many consensus heptads can be deleterious, even at CTD lengths comparable to the wild-type length.

### The CTD Dynamically Enters Sites of Active Transcription Independent of the Body of Pol II

Results from previous studies have lead to conjecture that transcription occurs in “transcription compartments” or “transcription factories” (Papantonis and Cook, 2013; Yao et al., 2007); more recent findings suggest that these could be phase-separated liquids formed by networks of weak interactions between RNA and proteins with multivalent, low complexity regions (Cho et al., 2018; Chong et al., 2018; Kwon et al., 2013; Lu et al., 2018; Sabari et al., 2018). In vitro, the CTD has been shown to partition into liquid and gel phases comprised of other low complexity domains and this partitioning is influenced by both the length and sequence composition of the CTD (Boehning et al., 2018; Kwon et al., 2013). Since transcriptionally active loci are easily observed as puffs on polytene chromosomes in *Drosophila* using microscopy, we explored whether the CTD serves to partition Pol II into sites of active transcription. To this end, we expressed a green fluorescent protein (GFP)-CTD fusion and a GFP control in salivary glands and analyzed the interaction of these proteins with polytene chromosomes. Each protein had a nuclear localization signal (NLS) and a FLAG-tag, and both were expressed at comparable levels and localized to nuclei (Figure S5). Indirect immunofluorescence microscopy of spread chromosomes revealed that GFPCTD co-localized with Pol II at puffs, sites of robust transcriptional activity, whereas GFP alone did not (Figure 4A and 4B). GFP-CTD was also enriched at puffs on polytene chromosomes in live cells whereas GFP was not (Figure 4C, red arrows). Thus, the CTD alone functions as a signal sequence that facilitates interaction of Pol II with transcriptionally active loci.

**Figure 4.**
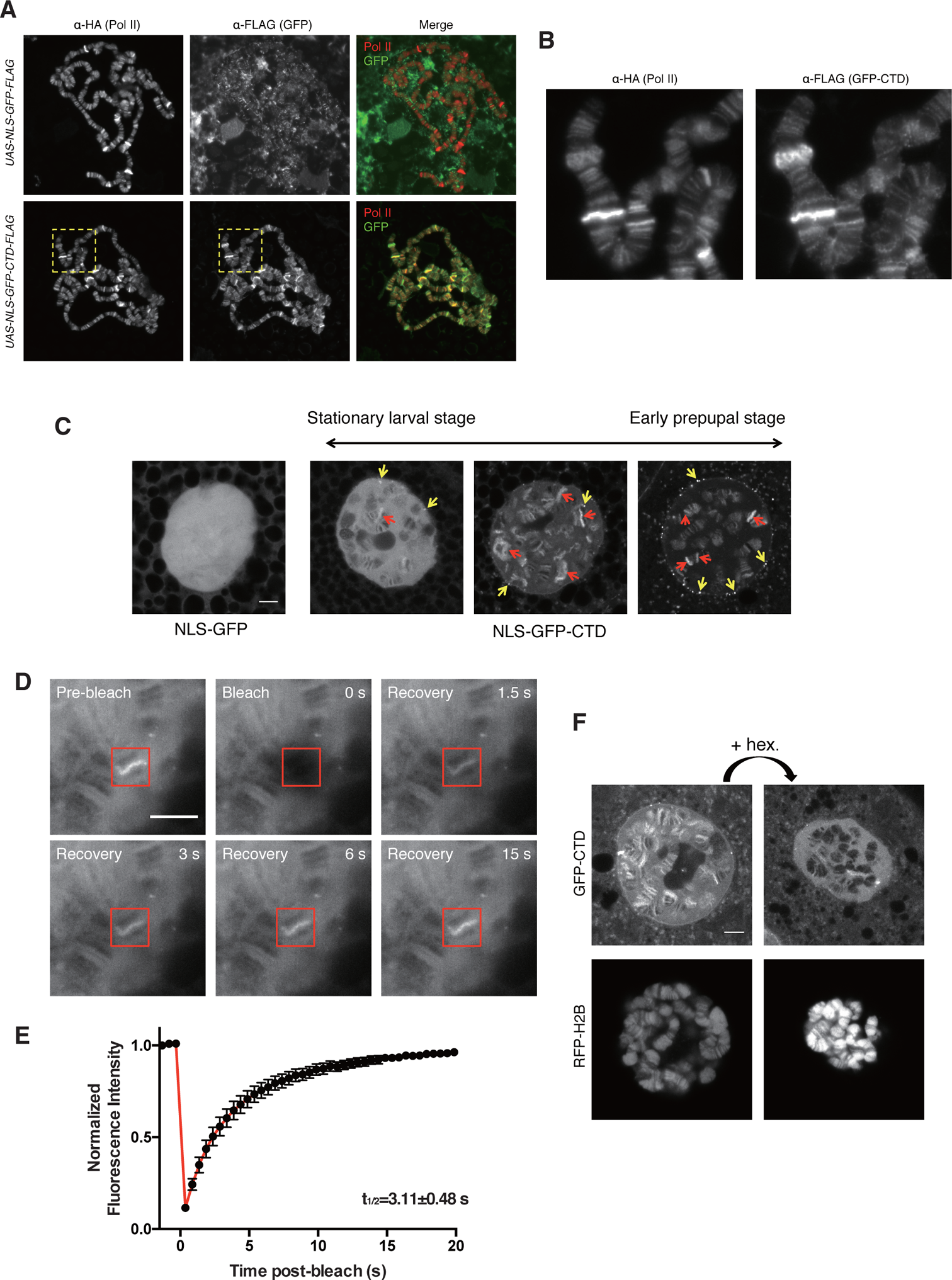
GFP-CTD is dynamically associated with sites of active transcription and sensitive to 1,6-hexanediol. (A) Indirect immunofluorescence detection of HA-tagged Pol II and FLAG-tagged NLSGFP derivatives on fixed and spread polytene chromosomes dissected from progeny derived from matings of *Rpb1_HA_*, *da-GAL4* to *UAS-NLS-GFP-CTD-FLAG* or *UAS-NLSGFP-FLAG* using anti-HA and anti-FLAG antibodies. See methods for details. (B) Magnified view of yellow boxes in (B). (C) Single-stack confocal images of nuclei in live salivary glands expressing either NLSGFP or NLS-GFP-CTD. >99% of cells expressing the NLS-GFP-CTD from the stationary larval stage have a range of appearances that resemble the left and middle panels whereas >99% cells from early prepupal stage have a range of appearances that resemble the middle and right panels. Examples represent the range of appearances observed. Red and yellow arrows indicate examples of chromosomal puffs and extrachromosomal foci that show enrichment of GFP-tagged CTD respectively. (D and E) FRAP images (D) and average GFP intensity at GFP-CTD puffs following photobleaching (E). The red box indicates a bleached region. Data are from 15 puffs in 15 different salivary glands. Error bars represents SEM. (F) Treatment with 1% 1,6-hexanediol disrupts the association of GFP-CTD with chromosomes. Fluorescence from mCherry-labeled H2B indicates that the chromosomes remain intact following 1,6-hexanediol treatment. Each image is a single Z stack. All scale bars represent 5 µm.

Higher resolution live cell imaging also revealed that GFP-CTD formed bright foci that were not on chromosomes whereas GFP alone did not (Figure 4C, yellow arrows and Figure S6). Fluorescence recovery after photobleaching (FRAP) showed that some GFP-CTD-containing foci recover while others do not (Figure S6, Movie S1 and S2), similar to recent findings for Pol II (Cho et al., 2018). Foci that did not recover from FRAP tend to be at the nuclear periphery separate from chromosomes. The bleached GFP-CTD recovers with a half time of approximately 3 seconds at positions on the chromosomes (Figure 4, D and E, Movie S3) and at a subset of foci (Figure S6, Movie S2), similar to the recovery rate observed with Pol II clusters (Cho et al., 2018).

The dynamic nature of the interaction of GFP-CTD with chromosomes and a subset of foci indicates the possibility that the CTD is partitioning into phase-separated liquid compartments. To test this, we examined the effect of treating glands with 1,6-hexanediol, which has been shown to disperse phase-separated compartments (Boehning et al., 2018; Cho et al., 2018; Kroschwald et al., 2017; Lu et al., 2018; Sabari et al., 2018; Strom et al., 2017). Treatment of intact glands with 1,6-hexanediol resulted in significant loss of GFP-CTD from chromosomes (Figure 4F, upper panels) and the disappearance of a subset of GFP-CTD foci (Figure S7). Notably, although the nucleus shrunk in size upon treatment with 1,6-hexanediol, the histone banding pattern was not disrupted (Figure 4F, lower panels), suggesting that the loss of GFP-CTD on chromosomes was not due to a global disruption of chromosome structure.

### Consensus CTDs at Wild-type Lengths Tend to Localize to Static Nuclear Foci Separate from Chromosomes

We next investigated the behavior of the consensus CTDs by fusing them to GFP and expressing them in salivary glands. Unlike GFP-CTD which is enriched at a few puffs during stationary larval stage and tends to be enriched at more puffs during early prepupal stage (Figure 4C), GFP-10con is not enriched on puffs and shows no increase in concentration on chromosomes over what is observed in the nucleoplasm (Figure 5A). Similar to GFP-CTD, GFP-20con, GFP-24con and GFP-29con can also be enriched on puffs, although the enrichment is not observed until early prepupal stage (Figure 5B and 5C, Figure S8A to S8D). This argues that the increase in CTD length can be more favorable for targeting to sites of active transcription.

**Figure 5.**
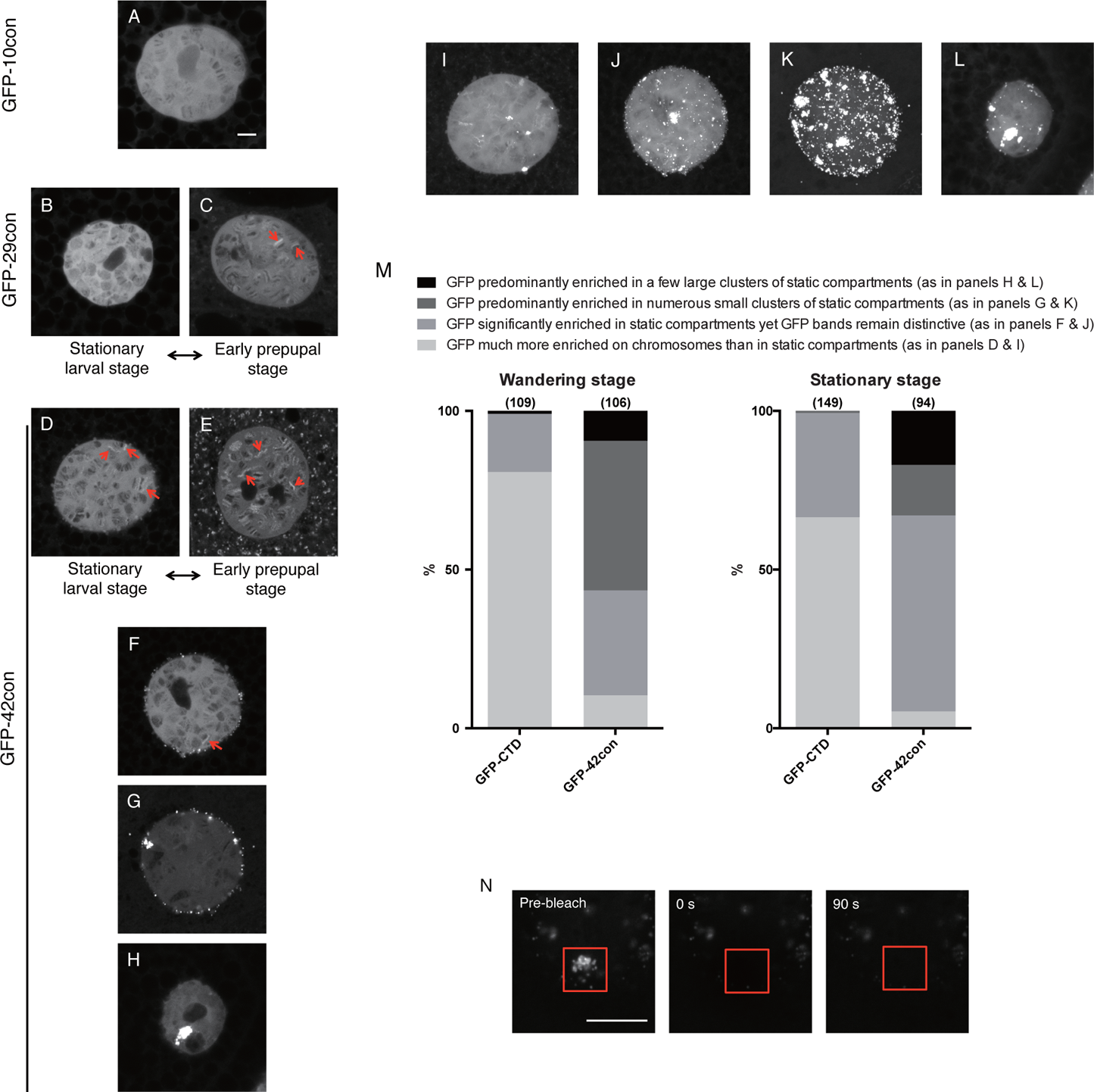
Varying the CTD sequence composition alters its distribution in the nucleus. (A to H) Single-stack confocal images of nuclei in live salivary glands expressing NLSGFP-10con (A), NLS-GFP-29con (B and C) and NLS-GFP-42con (D to H). B and D were collected from stationary larval stage whereas C and E were collected from early prepupal stage. Examples depicted in A, F, G and H can be observed in both stages. Red arrows indicate bright puffs that show GFP enrichment over nucleoplasm. (I to L) Maximum Intensity Projections of the same nuclei shown in D, F, G and H. (M) GFP-42con cells are more prone to partition in extrachromosomal stationary foci, much of which do not recover after photobleaching as shown in (N). Sample sizes are indicated in parentheses. The different shades of gray indicate phenotypes described above the graph. (N) FRAP images on a cluster of stationary GFP-42con foci. Red box indicates bleached region. All scale bars represent 5 µm.

The enrichment of the CTD on puffs can be observed in a subset of cells expressing either GFP-42con or GFP-52con at both stationary larval and early prepupal stages (Figure 5D and 5E, Figure S8E and S8F). However, GFP-42con and GFP-52con are more frequently localized in stationary foci than on chromosomes in the majority of cells at both stages (Figure 5D to 5M, Figure S8E to S8N). Much of the stationary foci containing GFP-42con do not recover after photobleaching (Figure 5N, Movie S4), suggesting that consensus CTDs at wild-type lengths are more prone to localize in static foci. Together, our results with GFP-CTD fusions supports the hypothesis that additional CTD length improves targeting to transcription compartments, but CTDs above a length threshold form less dynamic, possibly aggregated structures. This propensity for aggregation is counterbalanced by the divergent motifs that predominate in metazoa.

## Discussion

Results from our functional analysis of the CTD in *Drosophila* challenge the longstanding proposition that features of the CTD unique to multicellular organisms are essential for development. We show that substituting fully-consensus CTDs of 20 to 29 repeats supports *Drosophila* viability, development and stress tolerance. However, substituting fully-consensus CTDs of wild-type number of repeats or longer (42 or 52 repeats) do not. Remarkably, this optimal range in number of consecutive consensus heptads that functions in *Drosophila* is similar to the number consecutive consensus and near consensus heptads found in a wide spectrum of eukaryotes spanning from yeast to plants (Yang and Stiller, 2014). We propose that this consecutive array of consensus heptads functions as a single unit, otherwise it is hard to account for why the number of consecutive consensus heptads would be conserved within a given taxa such as chordates. Moreover, CTD mutant strains of yeast spontaneously acquire mutations that lead to CTDs consisting of approximately the natural number of 26 heptads (Morrill et al., 2016; Nonet and Young, 1989). How this consecutive array of repeats functions as a single unit remains to be determined but one possibility is that it provides for phosphorylation-dependent, highly cooperative regulation of factor binding (Lenz and Swain, 2006). This entropically driven mechanism of regulation could provide switch-like binding and release of factors dictated by threshold levels of phosphorylation.

To gain insight into what constraints impact the sequence composition of the CTD, we analyzed the targeting activity of various GFP-tagged CTD derivatives in *Drosophila* salivary glands where chromosomes are readily visible. A comparison of the targeting behavior of the 42 and 52 consensus CTDs, which do not fully support Pol II function, to those CTDs that do fully support CTD function indicates that divergent motifs could serve to counteract the propensity of the CTD to divert Pol II away from chromosomes. We observe that the *Drosophila* CTD and CTDs of 20, 24, and 29 consensus heptads associate with chromosomal puffs, which are well established to be regions of active transcription (Yao et al., 2008). In contrast to the fully function CTDs, the 42 and 52 consensus CTDs are less efficient at targeting puffs and instead associate more with static foci. We also frequently observe cases where the GFP-tagged 42 and 52 consensus CTDs form one or a few large clumps in the nucleus, a situation that rarely occurs with the fully functional CTDs. Our FRAP analysis reveals that the CTD interactions with puffs are dynamic whereas the interactions with many foci and large clumps are static. Recent studies show that isolated human and yeast CTDs self-associate into liquid phase separated droplets (Boehning et al., 2018). The low complexity, multivalent composition of the CTD probably drives this self-association. This propensity for self-association might be important for transcription since Pol II forms transient clusters whose duration and location correlate with transcriptional output (Cho et al., 2016). In cells, clustering and chromosome association of Pol II scaled with the number of heptad repeats that were attached to Pol II (Boehning et al., 2018). However, chromosome association was inferred from single molecule tracking rather than direct visualization of chromosomes. Given the 42 and 52 consensus CTDs are comparable in length to the *Drosophila* CTD and that the static structures formed by these consensus CTDs are clearly separate from chromosomes, we propose that divergent heptads function to counteract the propensity of extended arrays of consensus heptads to coalesce into dysfunctional states. This is consistent with the known behaviour of multivalent polymers where expansion of repeat numbers can cause molecules to transition from being dynamic to forming static, dysfunctional aggregates (Weber and Brangwynne, 2012).

In addition to allowing for functional self-association, the sequence composition of the CTD could also be constrained by its partition coefficient for liquid phase separated domains formed by other proteins. It was recently proposed that the puffs on polytene chromosomes could be liquid phase separated compartments (Harlen and Churchman, 2017). Our result showing that treatment of glands with 1,6-hexanediol caused the GFP-tagged CTD to disperse from puffs provides support for this proposition, since this dispersal is considered diagnostic of proteins partitioning into liquid phase separated domains (Boehning et al., 2018; Chong et al., 2018; Lu et al., 2018; Sabari et al., 2018). The failure of the GFP-tagged 10 consensus heptad to associate with puffs could be due to insufficient valency required for it to preferentially partition into puffs (Weber and Brangwynne, 2012).

We show that the malleability of the sequence composition of the *Drosophila* CTD is in stark contrast to the high level of sequence conservation displayed by members of the genus. We have identified targeting as a constraint to be added to the requirement for binding sites for CTD interacting factors. Consensus heptads could represent the ideal binding platform for both CTD-interacting factors and the interactions that facilitate CTD-self assembly in cells, but that too many such repeats results in aberrant dynamics with CTD-interacting factors and aberrant phase separation, resulting in the static foci observed in our studies. Substitution of consensus heptads with divergent motifs could provide a means to balance cooperative interactions that affect the presentation of binding sites in the CTD with phase separation.

## Acknowledgments

We thank M. M. Rolls, C. M. Thomas, and Z. C. Lai for imaging support; C. Feng for CRISPR and live imaging suggestions; D. G. Baumann, Y. Qiu, R. Dollinger and J. C. Reese for comments on the manuscript. The *Sgs3-GAL4, UAS-mCh-H2B/TM6* fly line is a generous gift from C. Peter Verrijzer. This work was funded by NIH GM047477 (to D.S.G.).

## Author Contributions

F.L. and D.S.G. planned the study and drafted the manuscript. B.P. designed and generated *UAS-NLS-GFP-CTD* fly line. D.S.G. generated *UAS-NLS-GFP* fly line and performed chromosome staining in Figure 4A and 4B. F.L. performed all other experiments. F.L., B.P. and D.S.G. revised the manuscript.

## Methods

Contact for Reagent and Resource Sharing

Further information and requests for resources and reagents should be directed to the Lead Contact, David S. Gilmour.

### Method Details

#### Sequence alignment

Previously published amino acid and nucleotide sequences of the CTDs of 12 species of *Drosophila* were downloaded from FLYBASE and aligned using Clustal Omega.

#### *Drosophila* strains

*UAS-Rpb1i, Act-GAL4* [*w; Act-GAL4* (*w+*)*/CyO*], *ms1096-GAL4* and *da-Gal4* fly lines were obtained from the Bloomington Stock Center (Bloomington 36830, 4414, 8860 and 55850 respectively). *UAS-Rpb1i*, *UAS-Rpb1^WT^* and *UAS-Rpb1i*, *UAS-Rpb1^human^* lines were previously described (Portz et al., 2017). DNA encoding either mutant Rpb1 with double FLAG-tags at the C-terminus, or various NLS-GFP derivatives were subcloned into the pUAST-attB vector, followed by transformation into the *PhiC31 attP 86Fb site* (Bischof et al., 2007). Rpb1i-resistance of transgenic Rpb1 was achieved by changing the part of the coding sequence of Rpb1 that is targeted by the 21 nucleotide shRNA (sense strand: AACGGTGAAACTGTCGAACAA) to AAC *C* GT *C* AA *GTTGAGC* AACAA. The *UAS-Rpb1i, UAS-Rpb1* line, as well as the w, *Rpb1_HA_*; *Act-GAL4*(*w+*) */CyO* and *Rpb1_HA_*; *da-GAL4* lines were generated by routine matings and meiotic recombination.

CRISPRed Rpb1 fly lines were generated through injection of DNA mixtures into a Cas9 expressing fly line (Bloomington 51324). As previously described (Gratz et al., 2014), each DNA mixture contains two CRISPR targeting vectors that express sgRNAs and a homology donor vector. The two CRISPR targets were selected from a list of targets with zero predicted potential off-target sites using flyCRISPR Optimal Target Finder (http://tools.flycrispr.molbio.wisc.edu/targetFinder/; with Maximum Stringency, NGG PAM sequence only). Complementary strands encoding the sgRNAs were annealed, phosphorylated and cloned into the BbsI site of pU6-BbsI-chiRNA (Addgene Plasmid 45946). The combination of Rpb1_fw1: 5’-CTTCGTAGGGATTTGAGAGCCAGTG-3’ and Rpb1_rev1: 5’-AAACCACTGGCTCTCAAATCCCTAC-3’ directed cleavage downstream of the 3’UTR of *Rpb1*. The combination of Rpb1_fw2: 5’-CTTCGAGAAACACTCGGCGAGGCT-3’and Rpb1_rev2: 5’-AAACAGCCTCGCCGAGTGTTTCTC-3’ directed cleavage adjacent to the start of the *Drosophila* CTD. Two homology arms, each about 1kb in length were PCR amplified from strain *w^1118^*. The homology arm adjacent to the 3’UTR of *Rpb1* was cloned into the AarI site of pHD-ScarlessDsRed (DGRC plasmid 1364). The other homology arm was cloned into the SapI site alongside desired mutations in the CTD and the CRISPR targeting site (Figure 2A). Candidate recombinant flies were identified by expression of the DsRed marker in eyes. Individual DsRed candidates were out-crossed to strain *w^1118^*. Desired candidates were verified by sequencing and western blot. The insertion of the DsRed cassette disrupts the gene adjacent to the 3’UTR of *Rpb1*, which to our knowledge does not affect the interpretation of our data. The Cas9 transgene on the third chromosome was subsequently removed by selection against the 3xP3-GFP marker. *Rpb1^42con^* and *Rpb1^52con^* alleles were maintained over *FM7(Tb)* balancer (BDSC 36337).

Illustration of genetic mating schemes in Figure 3A was generated using Genotype Builder (Roote and Prokop, 2013).

#### PCR genotyping

DNA was extracted from male individuals of desired genotypes as previously described (Gloor et al., 1993), and analyzed with the following primers: fw: 5’-CGCCTTCGGCTGCATCGG-3’ and rev: 5’-ACAAAGATCCTCTAGAGGTACCCTCGAGC-3’ (Figure S2); fw: 5’-GGCGATCGAGCGTAGTCGGTACTT-3’ and rev: 5’-CCAGGACCTTCGATGTCGCCGTATTT-3’ (Figure S4).

#### Western blotting

Lysate was prepared from late pupae (Figure 3, C and F) as previously described (Gibbs et al., 2017). Samples were subsequently loaded onto either a 7% Tris-Acetate gel (Figure 3, C and F). Rpb1_HA_ was detected with mouse anti-HA antibody monoclonal antibody (1:2500, Pierce, 26183, Thermo Scientific). Spt5 and Rpb3 were detected with rabbit anti-Spt5 antibody (1:3000) and rabbit anti-Rpb3 antibody (1:3000) respectively (Qiu and Gilmour, 2017). Expressions of FLAG Rpb1 and FLAG-GFP derivatives were detected with mouse anti-FLAG M2 antibody (1:2500; Sigma). Blots were subsequently probed with goat anti-rabbit IgG (1:3000; Alexa Fluor 488) and goat anti-mouse IgG (1:3000; Alexa Fluor 647) and visualized with a Typhoon (GE Healthcare).

#### Immunofluorescence

For polytene chromosome staining, salivary glands from wandering third-instar larvae were dissected and squashed as previously described (Schwartz et al., 2004). Slides were then incubated with rabbit anti-HA antibody (1:100; ThermoFisher 71-5500) and mouse anti-FLAG M2 antibody (1:200; Sigma) overnight at 4^°^C. For Figure 2G, the slides were subsequently probed with goat anti-rabbit IgG (1:200; Alexa Fluor 488) and goat anti-mouse IgG (1:200; Alexa Fluor 568) for 4 hours at room temperature. For Figure 4A and 4B, the slides were subsequently probed with goat anti-mouse IgG (1:200; Alexa Fluor 488) and goat anti-rabbit IgG (1:200; Alexa Fluor 568) for 4 hours at room temperature. Samples were imaged on a CARV II spinning disc confocal (BD Biosystems) and adjusted for brightness and contrast using ImageJ.

#### Animal assays

For Figure 2, D and E, animals were cultured at 30^°^C and 33% humidity. For Figure 3D, animals were cultured at 18^°^C and 33% humidity. Wings were dissected from 1 to 3 day old females. For cold shock analysis in Figure 2F, 0-6h old adults were aged for 96h, placed on ice for 14h and recover at room temperature (21^°^C). Recovery time was defined as the time point at which an individual started to move. For the rest of the figures, animals were maintained at 24^°^C and 65% humidity.

#### *Drosophila* live imaging

Larvae were raised in a non-crowded vial with standard cornmeal culture. Prior to dissection, wandering third instar larvae expressing a GFP-tagged CTD derivative fusion and mCherry tagged H2B-CTD (*Sgs3-GAL4, UAS-mCh-H2B/UAS-NLS-GFPCTD*) were transferred to a fresh vial. The larvae that were selected tended to migrate away from culture and always developed into early prepupae within 1 to 6 hours if left undissected, which corresponded to the “stationary larvae stage” (Fletcher and Thummel, 1995). Newly formed prepupae (<1 hour post puparium formation) were selected as the “early prepupae stage”. Individuals at this stage have transparent glands that can be clearly distinguished from prepupae of later stages, which have white non-transparent glands. Glands were dissected and imaged in M3 medium (Sigma Aldrich S8398) supplemented with 2.5 g of bactopeptone and 1 g of yeast extract per liter medium (M3+BPYE). Confocal images were acquired using a Zeiss LSM800 with a 63X oil objective (NA1.4) and adjusted for brightness and contrast using ImageJ. For consistency in comparison, all images of single nuclei were collected from the distal ⅓ of the salivary glands. Polytene nuclei were imaged at a depth no greater than 60 µm into the salivary gland tissue. For Figure 4C, Figure 5 and Figure S8, the intensity of GFP was adjusted based on the intensity of mCherry-H2B to correct for minor intensity differences produced by variation in imaging depth. Non-confocal images in Figure S5 were collected and adjusted with the same setting on a Carl Zeiss Axioskop 40 microscope.

#### Fluorescence recovery after photobleaching (FRAP)

Salivary glands were prepared as above. Three images were pre-bleach. On the fourth frame, a square region of approximately 4 µm X 4 µm was bleached with full laser power and recovery was recorded at 0.2% power, at 0.5 second intervals for 90 seconds. Normalized fluorescence intensity was calculated with a two-step normalization: first by normalizing the fluorescence intensity of GFP-CTD puffs to that of pre-bleach, second by normalizing this ratio to the ratio obtained from an unbleached region to correct for the subtle loss in fluorescence intensity due to bleaching. For calculating half time recoveries, normalized values from each recording were separately fit to a single exponential model, and half time recovery was presented as mean ± standard error.

#### 1,6-Hexanediol treatments

Salivary glands were prepared as above and immediately imaged in M3+BPYE (before treatment). Salivary glands were then transferred to the same medium containing 1% 1,6-hexanediol (Alfa Aesar A12439), incubated for approximately 20 min and imaged again (after treatment).

## Supplemental Information

Figure S1. Sequence alignments of the CTDs of *Drosophila*, related to Figure 1B.

(A) Nucleotide sequence alignment of the CTDs from 12 *Drosophila* species. The colors identify nucleotides that are identical (green) or different (magenta) compared with *D. melanogaster*.

(B) Amino acid sequence alignment of the CTDs from 12 *Drosophila* species. The colors identify amino acids that are identical (green), similar (yellow) or dissimilar (magenta) compared with *D. melanogaster*. Pink boxes highlight the only two consensus heptads in the CTD of *D. melanogaster*.

Figure S2. PCR validation of various fly lines carrying *UAS-Rpb1* transgenes, related to Figure 3C. One primer was specific to *UAS-Rpb1* insertion and the other hybridized to the body of *Rpb1*.

Figure S3. Replacing the endogenous Rpb1 with either Rpb1^20con^ or Rpb1^24con^ using CRISPR-Cas9 produces healthy homozygous flies, related to Figure 3E.

(A) Hatch rates of embryos at 24^°^C, measured 36h after egg deposition, n>350 for each genotype.

(B) Percentages of hatched embryos that developed into adults when raised at 24^°^C, n>300 for each genotype.

(C) Hatch rates of embryos at 30^°^C, measured 36h after egg deposition, n>350 for each genotype.

(D) Percentages of hatched embryos that developed into pupae when raised at 30^°^C, n>400 for each genotype.

(E) Quantification of recovery time after 14h of cold shock on ice, n=64 for *Rpb1_FLAG_*; n=64 for *Rpb1^20con^*; n=68 for *Rpb1^24con^* (equal numbers of male and female adults were assayed). Dots represent individual adults. Red bars show the medians and interquartile ranges.

Figure S4. PCR validation of hemizygotic males with the modified endogenous Rpb1 locus using primers flanking the CTD, related to Figure 3F. A male fly with unmodified Rpb1 locus [*w*(control)] provides the PCR product of the unmodified locus. Changes in the sizes of the PCR products reflect modification of the endogenous Rpb1 locus.

Figure S5. Fluorescence detection of NLS-GFP derivatives in the nuclei of salivary glands, related to Figure 4 and Figure 5. Expression was driven by *Sgs3-GAL4*.

Figure S6. FRAP analysis reveals that GFP-CTD containing foci consist of static (upper) and dynamic (lower) compartments, related to Figure 4. Scale bar represents 5 µm.

Figure S7. Treatment with 1% 1,6-hexanediol dissolved a subset of GFP-CTD foci, related to Figure 4F. Maximum intensity projections of entire Z stacks. Scale bar represents 5 µm. The diffuse fluorescence detected after 1,6-hexanediol treatment is due to the release of GFP-CTD from the chromosomes.

Figure S8. Varying the CTD sequence composition alters its distribution in the nucleus, related to Figure 5.

(A to I) Single-stack confocal images of nuclei in live salivary glands expressing NLSGFP-20con (A and B), NLS-GFP-24con (C and D) and NLS-GFP-52con (E to I). A, C and E were collected from stationary larval stage whereas B, D and F were collected from early prepupal stage. Examples depicted in G, H and I can be observed in both stages. Scale bar represents 5 µm. Red arrows demarcate examples of puffs that show enrichment over the nucleoplasm.

(J to M) Maximum Intensity Projections of the same nuclei shown in E, G, H and I, respectively.

(N) GFP-52con cells are more prone to partition in static compartments. Sample sizes are indicated in parentheses.

Movie. S1. FRAP movie on static GFP-CTD foci, related to Figure S6 (upper).

Movie. S2. FRAP movie on dynamic GFP-CTD foci, related to Figure S6 (lower).

Movie. S3. Representative FRAP movie on a GFP-CTD puff, related to Figure 4D.

Movie. S4. Example of stationary foci containing GFP-42con that did not recover after photobleaching, related to Figure 5N.

## References and Notes

Bischof, J., Maeda, R.K., Hediger, M., Karch, F., and Basler, K. (2007). An optimized transgenesis system for Drosophila using germ-line-specific φC31 integrases. Proc. Natl. Acad. Sci. U. S. A. 104, 3312–3317.

Boehning, M., Dugast-Darzacq, C., Rankovic, M., Hansen, A.S., Yu, T., Marie-Nelly, H., McSwiggen, D.T., Kokic, G., Dailey, G.M., Cramer, P., et al. (2018). RNA polymerase II clustering through carboxy-terminal domain phase separation. Nat. Struct. Mol. Biol.

Chapman, R.D., Heidemann, M., Hintermair, C., and Eick, D. (2008). Molecular evolution of the RNA polymerase II CTD. Trends Genet. 24, 289–296.

Cho, W.-K., Jayanth, N., English, B.P., Inoue, T., Andrews, J.O., Conway, W., Grimm, J.B., Spille, J.-H., Lavis, L.D., Lionnet, T., et al. (2016). RNA Polymerase II cluster dynamics predict mRNA output in living cells. Elife 5.

Cho, W.-K., Spille, J.-H., Hecht, M., Lee, C., Li, C., Grube, V., and Cisse, I.I. (2018). Mediator and RNA polymerase II clusters associate in transcription-dependent condensates. Science.

Chong, S., Dugast-Darzacq, C., Liu, Z., Dong, P., Dailey, G.M., Cattoglio, C., Heckert, A., Banala, S., Lavis, L., Darzacq, X., et al. (2018). Imaging dynamic and selective low-complexity domain interactions that control gene transcription. Science eaar2555.

Corden, J.L. (2013). RNA polymerase II C-terminal domain: Tethering transcription to transcript and template. Chem. Rev. 113, 8423–8455.

Dias, J.D., Rito, T., Torlai Triglia, E., Kukalev, A., Ferrai, C., Chotalia, M., Brookes, E., Kimura, H., and Pombo, A. (2015). Methylation of RNA polymerase II non-consensus Lysine residues marks early transcription in mammalian cells. Elife 4.

Egloff, S., Dienstbier, M., and Murphy, S. (2012). Updating the RNA polymerase CTD code: adding gene-specific layers. Trends Genet. 28, 333–341.

Eick, D., and Geyer, M. (2013). The RNA polymerase II carboxy-terminal domain (CTD) code. Chem. Rev. 113, 8456–8490.

Fletcher, J.C., and Thummel, C.S. (1995). The Drosophila E74 gene is required for the proper stage-and tissue-specific transcription of ecdysone-regulated genes at the onset of metamorphosis. Development 121, 1411–1421.

Gibbs, E.B., Lu, F., Portz, B., Fisher, M.J., Medellin, B.P., Laremore, T.N., Zhang, Y.J., Gilmour, D.S., and Showalter, S.A. (2017). Phosphorylation induces sequence-specific conformational switches in the RNA polymerase II C-terminal domain. Nat. Commun. 8, 15233.

Gloor, G.B., Preston, C.R., Johnson-Schlitz, D.M., Nassif, N.A., Phillis, R.W., Benz, W.K., Robertson, H.M., and Engels, W.R. (1993). Type I repressors of P element mobility. Genetics 135, 81–95.

Gratz, S.J., Ukken, F.P., Rubinstein, C.D., Thiede, G., Donohue, L.K., Cummings, A.M., and O’Connor-Giles, K.M. (2014). Highly specific and efficient CRISPR/Cas9-catalyzed homology-directed repair in Drosophila. Genetics 196, 961–971.

Harlen, K.M., and Churchman, L.S. (2017). The code and beyond: transcription regulation by the RNA polymerase II carboxy-terminal domain. Nat. Rev. Mol. Cell Biol. 18, 263–273.

Kroschwald, S., Maharana, S., and Simon, A. (2017). Hexanediol: a chemical probe to investigate the material properties of membrane-less compartments. Matters 3, e201702000010.

Kwon, I., Kato, M., Xiang, S., Wu, L., Theodoropoulos, P., Mirzaei, H., Han, T., Xie, S., Corden, J.L., and McKnight, S.L. (2013). Phosphorylation-regulated binding of RNA polymerase II to fibrous polymers of low-complexity domains. Cell 155, 1049–1060.

Lenz, P., and Swain, P.S. (2006). An entropic mechanism to generate highly cooperative and specific binding from protein phosphorylations. Curr. Biol. 16, 2150–2155.

Liu, P., Kenney, J.M., Stiller, J.W., and Greenleaf, A.L. (2010). Genetic organization, length conservation, and evolution of RNA polymerase II carboxyl-terminal domain. Mol. Biol. Evol. 27, 2628–2641.

Lu, H., Yu, D., Hansen, A.S., Ganguly, S., Liu, R., Heckert, A., Darzacq, X., and Zhou, Q. (2018). Phase-separation mechanism for C-terminal hyperphosphorylation of RNA polymerase II. Nature 558, 318–323.

Morrill, S.A., Exner, A.E., Babokhov, M., Reinfeld, B.I., and Fuchs, S.M. (2016). DNA Instability Maintains the Repeat Length of the Yeast RNA Polymerase II C-terminal Domain. J. Biol. Chem. 291, 11540–11550.

Nonet, M.L., and Young, R.A. (1989). Intragenic and extragenic suppressors of mutations in the heptapeptide repeat domain of Saccharomyces cerevisiae RNA polymerase II. Genetics 123, 715–724.

Papantonis, A., and Cook, P.R. (2013). Transcription factories: genome organization and gene regulation. Chem. Rev. 113, 8683–8705.

Portz, B., Lu, F., Gibbs, E.B., Mayfield, J.E., Rachel Mehaffey, M., Zhang, Y.J., Brodbelt, J.S., Showalter, S.A., and Gilmour, D.S. (2017). Structural heterogeneity in the intrinsically disordered RNA polymerase II C-terminal domain. Nat. Commun. 8, 15231.

Qiu, Y., and Gilmour, D.S. (2017). Identification of regions in the Spt5 subunit of DSIF that are involved in promoter proximal pausing. J. Biol. Chem.

Roote, J., and Prokop, A. (2013). How to design a genetic mating scheme: a basic training package for Drosophila genetics. G3 3, 353–358.

Sabari, B.R., Dall’Agnese, A., Boija, A., Klein, I.A., Coffey, E.L., Shrinivas, K., Abraham, B.J., Hannett, N.M., Zamudio, A.V., Manteiga, J.C., et al. (2018). Coactivator condensation at super-enhancers links phase separation and gene control. Science.

Schröder, S., Herker, E., Itzen, F., He, D., Thomas, S., Gilchrist, D.A., Kaehlcke, K., Cho, S., Pollard, K.S., Capra, J.A., et al. (2013). Acetylation of RNA polymerase II regulates growth-factor-induced gene transcription in mammalian cells. Mol. Cell 52, 314–324.

Schwartz, B.E., Werner, J.K., and Lis, J.T. (2004). Indirect immunofluorescent labeling of Drosophila polytene chromosomes: visualizing protein interactions with chromatin in vivo. Methods Enzymol. 376, 393–404.

Simonti, C.N., Pollard, K.S., Schröder, S., He, D., Bruneau, B.G., Ott, M., and Capra, J.A. (2015). Evolution of lysine acetylation in the RNA polymerase II C-terminal domain. BMC Evol. Biol. 15, 35.

Sims, R.J., Rojas, L.A., Beck, D.B., Bonasio, R., Schüller, R., Drury, W.J., 3rd, Eick, D., and Reinberg, D. (2011). The C-terminal domain of RNA polymerase II is modified by site-specific methylation. Science 332, 99–103.

Strom, A.R., Emelyanov, A.V., Mir, M.R., Fyodorov, D.V., Darzacq, X.R., and Karpen, G.H. (2017). Phase Separation Drives Heterochromatin Domain Formation. Nature 547, 241–245.

Weber, S.C., and Brangwynne, C.P. (2012). Getting RNA and protein in phase. Cell 149, 1188–1191.

Yang, C., and Stiller, J.W. (2014). Evolutionary diversity and taxon-specific modifications of the RNA polymerase II C-terminal domain. Proc. Natl. Acad. Sci. U. S. A. 111, 5920–5925.

Yao, J., Ardehali, M.B., Fecko, C.J., Webb, W.W., and Lis, J.T. (2007). Intranuclear distribution and local dynamics of RNA polymerase II during transcription activation. Mol. Cell 28, 978–990.

Yao, J., Zobeck, K.L., Lis, J.T., and Webb, W.W. (2008). Imaging transcription dynamics at endogenous genes in living Drosophila tissues. Methods 45, 233–241.

Zaborowska, J., Egloff, S., and Murphy, S. (2016). The pol II CTD: new twists in the tail. Nat. Struct. Mol. Biol. 23, 771–777.

Zhao, D.Y., Gish, G., Braunschweig, U., Li, Y., Ni, Z., Schmitges, F.W., Zhong, G., Liu, K., Li, W., Moffat, J., et al. (2016). SMN and symmetric arginine dimethylation of RNA polymerase II C-terminal domain control termination. Nature 529, 48–53.

